# Towards a highly efficient diversity census of the prokaryotic biosphere: a group testing approach

**DOI:** 10.1101/167502

**Authors:** Bar Shalem, Amnon Amir, Ely Porat, Noam Shental

## Abstract

Exploring the microbial biosphere has grown exponentially in recent years, although we are far from understanding its entirety. We present the âdiversity censusâ problem of exploring all bacterial species in a large cohort of specimens, and detecting a specimen that contains each species. The naive approach to this problem is to sequence each specimen, thus requiring costly sample preparation steps.

We suggest an orders of magnitude more efficient approach for diversity censusing. Specimens are pooled according to a predefined design and standard 16S rRNA sequencing is performed over each pool. For each bacterial species, from the ultra-rare to the most common, the algorithm detects a single specimen that contains the bacterial species. The approach can be applied to large cohorts of monomicrobial cultures or to complex samples containing a mixture of organisms.

We model the experimental procedure and show via *in silico* simulations that the approach enables censusing more than 95% of the species while taking 10 – 70 fold less resources. Simulating experiments using real samples display the utility in censusing large cohorts of samples.

Diversity censusing presents a novel problem in the mathematical field of group testing that may also be applied in other biological problems and in other domains.

## 1 Introduction

Technological advancements in recent years have elucidated the extent of the microbial “dark matter”. Replacing culture-based profiling by Next Generation Sequencing (NGS) methods paved the way to exploring the microbial content in different niches. However, we are far from capturing the microbial diversity across the prokaryotic biosphere, whose estimated size varies between 10^6^ species [1] to an incredible number of 10^12^ species [2]. Currently known species are highly biased towards host-associated niches while soil and aquatic environments, that are known for their high diversity and richness, take less than 25% of known species [3]. Moreover, even in well explored niches such as the human gut, known species are mainly biased towards the common ones while rare species remain unknown.

The objective of a “diversity census” is to enrich our knowledge of organisms and close the current gap. Organisms may be recorded based on their 16S rRNA gene, or, preferably, by their whole genome sequence. Moreover, once a novel organism is identified it may also be beneficial to actually culture it. Hence, a second census objective may be to create a repository of most organisms on Earth, both common and rare. Such a repository may have a tremendous impact since newly identified species may be candidates for new therapeutics, biotechnological applications, as well as important for basic research.

The first step in creating a census is to collect a large number of specimens. These may require global efforts, *e.g*., the Earth Microbiome Project [4], although the same questions may also be relevant for more ‘local’ projects. For example, a researcher may collect many specimens from a certain region in the ocean or in a desert aiming to create a diversity census.

The second step in a census is to list all organisms present in the set of specimens and detect at least one specimen that includes each species. The naive approach to censusing is to profile each specimen individually, as performed in all current large scale initiatives (*e.g*., National Microbiome Initiative [5], Earth Microbiome Project [4], Extreme Microbiome Project *etc*.). This approach results in a large amount of information which is not required for diversity censusing (*i.e*.the identity and frequency of all species in each specimen), while bearing large costs of processing all specimens and thus limiting the number specimens that may be screened.

The third potential step in creating a census is to culture organisms from specific specimens. As is often the case in many microbiome studies, it is desirable to return to a tested specimen and culture specific bacteria for downstream scientific research or for biotechnological applications (*e.g*., finding the ‘most wanted’ taxa from the Human Microbiome Project [6]). Although culturing bears great complications (classical culturing is able to grow only a small fraction of the microbial world) significant progress in this field of culturing techniques, often referred to as CulturOMICS [7, 8], has increased success rate dramatically. CulturOMICS applies miniaturization and compartmentalization approaches (*e.g*. via microdroplets and microfluidics [9, 10]) together with novel physico-chemical culture conditions that allow culturing of bacteria that were considered uncultivable in classical Petri dishes. Basically, the procedure involves isolating cells and culturing them, but since bacterial identity can not be deciphered *a priori* this procedure is repeated many times, especially if the abundance of bacteria on the ‘most wanted list’ is low. The latter results in huge repositories (> 10, 000 cultures) that need to be screened in order to detect a single culture for each species.

In this work we replace the naive ‘blind’ censusing by an approach much more suitable for a diversity census. For each bacterial species (from the ultra-rare to the highly common across the cohort of specimens) this highly efficient approach detects a *single* specimen containing that species.

In the context of this manuscript the word “specimens” refers to both monomicrobial *cultures* and to *samples* that contain a mixture of bacterial species. For practical reasons we distinguish between these two scenarios: (A) For each species in a large set of monomicrobial *cultures* the approach efficiently finds a *single* culture that contains it (Fig 1A); (B) Given a large set of *samples*, each containing a mixture of species, the approach efficiently finds a *single* sample that contains each bacterial species (Fig 1B). In this case we aim to find a sample in which the abundance of the relevant species is high enough. Upon detecting a single sample for each species, culturing may be applied to recover the bacterium. The difference between settings (A) and (B) stems from the fact that each culture in (A) contains a single bacterium, as opposed to multiple bacterial species in (B). Prior to describing our method we describe the current approach for diversity censusing.

**Figure 1:**
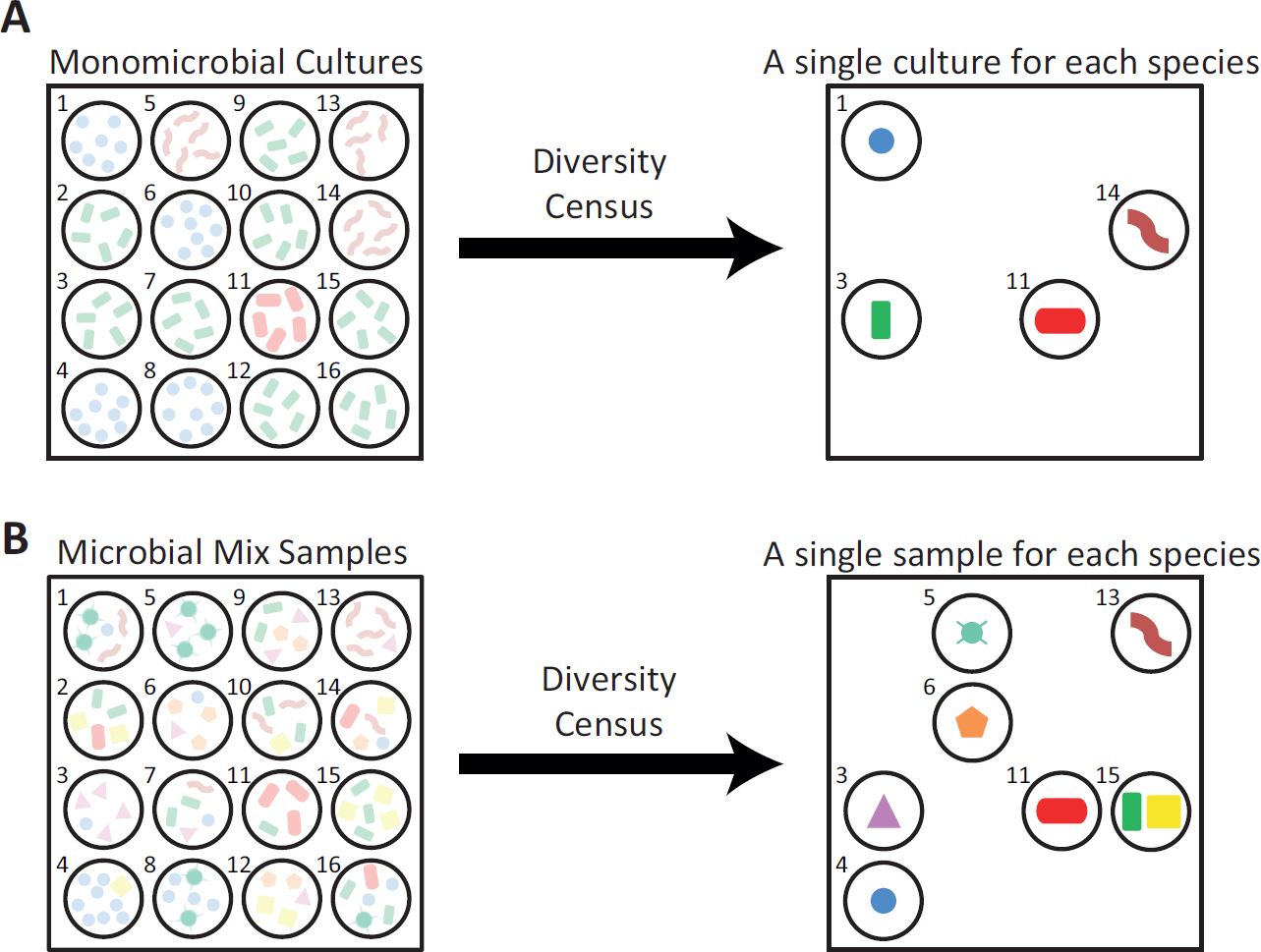
A bacterial diversity census. The left column presents cartoons of the bacterial identities in a cohort of 16 monomicrobial cultures (upper subplot) or samples (lower subplot). This ground truth (shown as opaque) is, of course, provided only for demonstration purposes, and is unknown in practice. The right column shows the required output of a diversity census of each cohort of specimens. A. Diversity census of a monomicrobial *culture* repository results in detecting a single culture that contains each bacterium in the cohort (depicted by the bacterial species in the center of each dish). Since the cohort contains four bacterial species diversity censusing should detect four cultures. Cultures that contain the same bacterium are ‘symmetric’ for the purpose of diversity censusing, *e.g*. each of the eight cultures that contain the ‘green-bacilli-like’ bacterium may be equally detected. B. Diversity censusing a repository of *samples*, where each sample contains many species, should detect a sample for each bacterial species across the cohort. For each bacterium, censusing should preferably detect a sample in which the bacterium abundance is high, and hence samples that contain the same bacterium are not ‘symmetric’. There are eight ‘species’ in this cartoon and seven detected samples, since two bacteria were detected in the same sample.

### Current work - the naive approach

The naive approach for diversity censusing is to screen each specimen individually, and thus obtain the content of each specimen. Screening a cohort of monomicrobial cultures or a set of complex samples (*i.e*., bacterial mixtures) is performed using different methods.

Screening a large set of *cultures*: The straightforward approach is to sequence each colony, either via Sanger sequencing or via barcoded multiplexed NGS, which involve costly and labor intensive sample preparation steps. Hence, screening large cohorts of *culture*s is not regularly performed by NGS but rather by Mass Spectrometry (MS) technology, or more often via matrix assisted laser desorption/ionization time-of-flight MS (MALDI-TOF). As opposed to NGS, sample preparation is not required and results are available within minutes at a negligible cost. Hence all cultures are processed and the identity of bacteria in a specific colony is deciphered by comparing its MS spectrum to a databases of known bacterial spectra. Since the seminal work of Seng *et al*. [11] MALDI-TOF has become a well established approach in clinical applications, although it may still be less efficient in research applications. The main shortcoming of MS is that its spectra database is still much smaller than the databases of millions of 16S rRNA gene sequences, and hence bacteria may be identified by NGS but not by MS. For a comprehensive and thorough review of MS and MALDI-TOF microbial identification see [12, 13] and references therein.

Screening a large set of *samples*: Following their collection, each sample is independently processed for 16S rRNA gene NGS sequencing (*i.e*., DNA is extracted, PCR amplified and sequenced). The composition of each sample is then discovered, and hence all samples containing a given bacterial sequence can be identified. However, this incurs the cost of processing and sequencing the full set of samples (currently, MALDI-TOF based identification can not be applied to a complex community of bacteria, *i.e*., to *samples* [14]).

### Our approach

We describe a generic approach for efficient diversity censusing of large repositories of specimens, *i.e., samples* or *cultures*, using sequencing of the 16S rRNA gene. The approach harnesses the highly accurate and comprehensive bacterial identification of NGS, while significantly reducing experimental costs.

The idea is to create *pools* of specimens according to a predefined design and then sequence them as if these were single specimens. Although information about the content of individual specimens is lost, the specific pooling design and the detection algorithm allow a diversity census, *i.e*., for each bacterial species a *single* specimen that contains each species is identified (bacterial species in this sense refer to both known and novel 16S rRNA gene sequences that appeared in any of the specimens, irrespective of their abundance across specimens). The major costs in NGS are related to pre-sequencing procedures (*i.e*. ‘sample preparation’), which in our approach are applied to the pools rather than to the specimens themselves. Hence, since the number pools is orders of magnitude lower than the number of specimens, censusing may be achieved with only a fraction of the costs compared to the naive approach and thus allow screening of larger cohorts. Diversity censusing via pooling is different from former pooled measurements approaches, as explained after describing the cases of cultures and samples.

Diversity censusing of a large cohort of *cultures*: For each bacterial species in a large repository of cultures, we identify a single culture that contain that species. Equal amounts of material from each culture are mixed into pools according to a predefined design, then DNA is extracted from each pool and standard 16S rRNA sequencing is performed over the pools as if they were a single culture. Based on profiling results and on the pooling design we then computationally detect a representative culture for each species. All steps are performed over the pools, whose number is about 50 fold smaller than the number of samples, which results in a dramatic reduction in costs. For example, the approach can detect *all* bacterial species of abundance 0–50% out of a set of 10, 000 samples with about 150 16S rRNA library preparations and NGS, namely the whole process requires 70 fold less resources (by abundance of 0% we refer to the most rare species in the set of cultures, *i.e*.,1/10, 000 in this example). The cartoon in Fig 2A displays censusing of four unidentified cultures (step (*i*)) resulting in identifying three bacterial species and a culture that contains each of them (step (*v*)). Censusing is performed by creating three pools comprised of taking material from specific cultures (*e.g*., in step (*ii*) the top pool is comprised of material from cultures *C*1, *C*2 and *C*4), and analyzing sequencing results (*e.g*., in step (*ii*) the top pool contains three species). We note in passing that due to the increased efficiency censusing does not provide the identity of all cultures (*e.g*., the third culture remains unknown).

**Figure 2:**
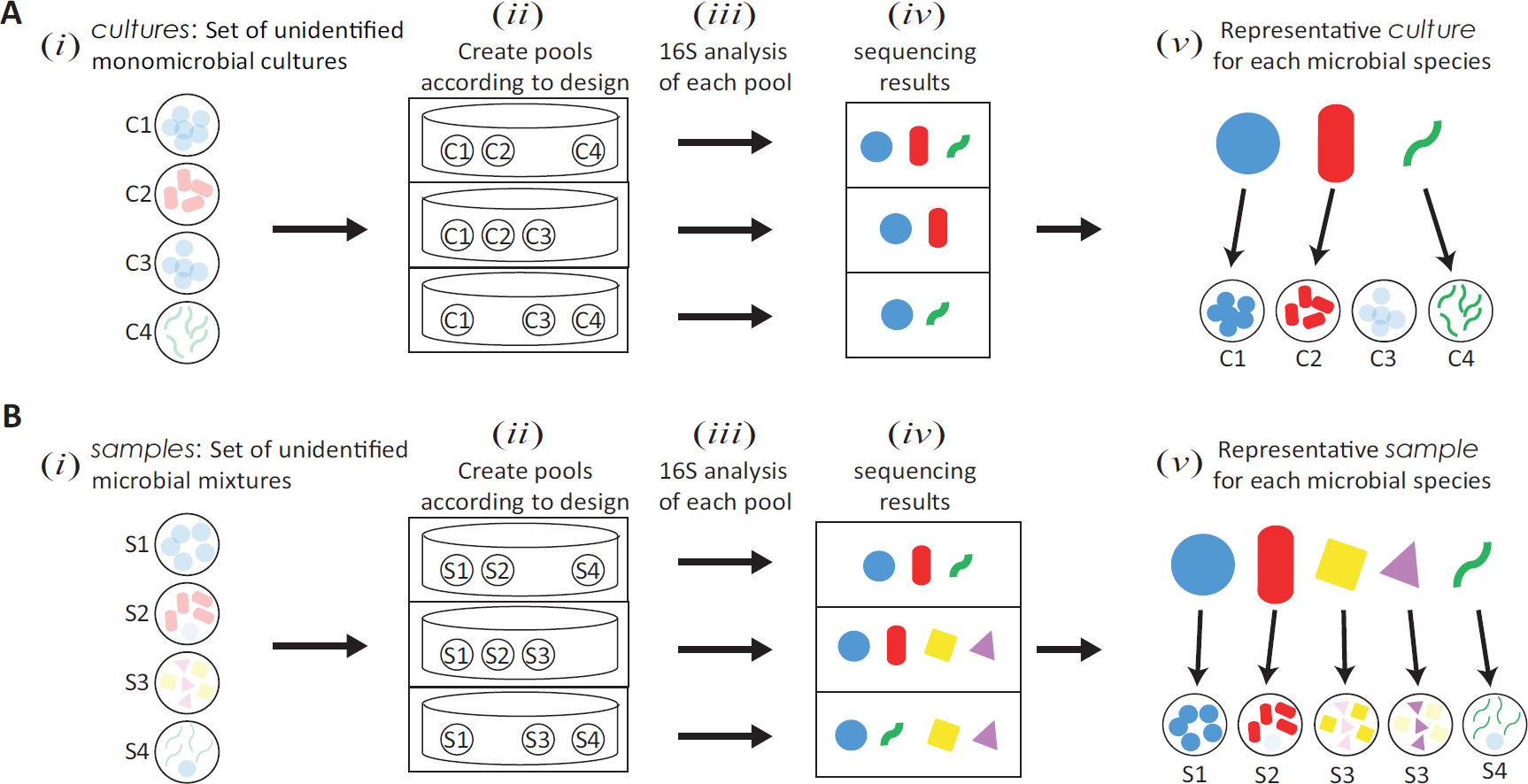
A diagram of the suggested approach. A. Diversity censusing *culture* repositories. The input is a set of cultures (*C*1 *–C*4), where each culture contains a single unknown species ((*i*), species in each culture appear as opaque to designate that their identities are unknown). Equal amounts of material from each culture are pooled according to a predefined design (*ii*), then DNA is extracted and 16S rRNA NGS is performed (*iii*). The sets of bacteria identified in each pool (*iv*) are analyzed to detect one culture for each bacterial species, independent of the their abundance (*v*). B. Diversity censusing repositories of *samples*, where each sample (*S*1 *–S*4) contains many species (*i*). The flowchart is similar to the case of cultures. The approach detects a representative sample for each bacterial species that appears in at least one sample and whose abundance is higher than some predefined threshold (*v*). The assigned sample to the ‘triangle’ and ‘square’ bacteria was the same one (*S*3).

Diversity censusing a large cohort of *samples*: For each bacterial species in a large repository of samples, we identify a single sample that contains that species, irrespective of its abundance across samples. Moreover, we aim to detect a sample in which the species’ abundance is high enough. A cartoon of the procedure is analogous to the case of cultures (Fig 2B). The input is a cohort of four samples, each containing many bacterial species (step (*i*)), and the output detects five bacterial species and samples that include each of them (step (*v*), sample *S*3 was detected twice in the context of two different species).

### Group testing

Methods for detecting specific specimens from a large cohort via pooled measurements are known as the mathematical field of group testing (GT), which was first applied by the US army during world war II to efficiently detect syphilitic draftees [15]. Since then the mathematical foundations of this field have been established and GT has been applied to other fields such as data compression [16], quality control in product testing [17], computation in the data stream model [18] and in screening large drug libraries. In former works we described a combination of GT and NGS for detecting carriers of rare genetic mutations[19, 20].

The canonical problem in GT aims to efficiently detect *rare* ‘faulty’ specimens out of a large cohort of ‘native’ specimens. Many algorithms and highly intricate pooling designs were presented in the last 70 years, in conjunction with theoretical analysis thus providing bounds on the required number of pools in various cases.

In this work we present a novel variation of the GT problem. As opposed to the classical GT problem which aims to locate *all* ‘faulty’ specimens and thus is restricted to detecting only *rare* specimens, we aim to detect a (single) *representative* of each ‘faulty’ type (namely species) while being independent of its abundance. Intuitively, pooled measurements are inherently limited in terms of the amount of information they provide regarding individual specimens, and thus a compromise regarding the required output has to be made. We simply relax one constraint of classical GT, and thus gain power where classical GT fails.

In this work we provide a pooling design and a detection algorithm that solves this variation of the GT problem highly efficiently. This somewhat heuristic design and algorithm calls for rigorous theoretical analysis of this novel GT problem which we hope will be followed by others in the future (theoretical bounds of a similar problem is a notoriously hard open question[21]). We refer to this GT-based diversity censusing approach as *GT-Census*.

### Outline

Details of the pooling design and the detection algorithm of GT-Census are provided in Materials and Methods. Results present the performance of GT-Census of a cohort of cultures when introducing different types of measurement errors. For the case of censusing cohorts of samples we provide simulations of pooling data from large scale experiments from the American Gut project [22] and from a longitudinal survey of humans and their indoor environments [23]. Future directions are presented in the Discussion.

## 2 MATERIALS AND METHODS

### Overview

The GT-Census approach, illustrated in Fig 2, is composed of the following steps.

i. *Input*: A large set of cultures or a large set of samples.
ii. *Pooling*: A pool is created by drawing equal amounts of material from each culture or sample according to a predefined design.
iii. *Sample preparation and sequencing*: DNA is extracted from each pool and 16S rRNA sequencing is performed, as if they were standard samples.
iv. *Microbial profiling*: A list of all bacteria in each pool is created. A bacterium is defined as ‘present’ in a pool if its corresponding number of reads is higher than some threshold, and ‘absent’ otherwise.
v. *Output*: A detection algorithm is independently used for each bacterial species that was present in any of the pools. For the case of *cultures* the algorithm outputs a single culture for each bacterium whose abundance in the repository is within the range of 0%-50%. For *samples*, the algorithm outputs a single sample for each bacterium whose inter-sample abundance is within the range of 0%-50%.

For simplicity we first consider the case of censusing a set of cultures. Following some definitions the detection algorithm in the absence of experimental noise is described. This algorithm provides an intuition for designing the measurement matrix outlined afterwards. We then describe the issue of experimental noise and the detection algorithms of cultures and of samples. Finally, the simulations performed to evaluate the performance of GT-Census are described.

### Definitions

The following definitions are used:

- The number of cultures in the repository is *N*. The number of pools is *T*.
- A‘*positive*’ or a ‘*negative*’ pool: A pool is defined as ‘positive for bacterium *r*’ if this bacterium was ‘present’ in that pool, or ‘negative for bacterium *r*’ if the bacterium was ‘absent’ from the pool. A pool is ‘positive for bacterium *r*’if at least one of its specimens contains *r*. Since identification is performed for each bacterium independently, we use the terms ‘positive’ or ‘negative’ without specifying the bacterium.
- Pooling design: the matrix *M*, that has *N* columns and *T* rows specifies the pooling design. An entry *M_ij_* is 1 if specimen *j* participates in pool *i* and 0 otherwise. The columns of *M* are unique, namely any pair of columns differ by at least one entry (*i.e*. pool).
- The measurement vector *y_r_* is a vector of length *T*, where *y_r_*(*i*)=1 if the *i′*th pool is positive for bacterium *r* and *y_r_*(*i*)=0 if pool *i* is negative for bacterium *r*.
- The set of specimens that contain bacterium *r* is *S_r_*. This set is, of course, unknown and our goal is to correctly identify one of its specimens.

We first consider the noiseless case where all pools are correctly called either as positive or as negative with respect to a bacterium *r*, and then discuss the case of measurement errors.

### Detection algorithm - the noiseless case

The following two-step algorithm is independently applied to each bacterium *r*, that appeared at least once across the pools, in order to detect an specimen that contains *r*.

- *Elimination step*: Since negative pools do not contain *r*, all specimens that belong to these pools may be deleted from the list of candidate specimens. Hence, only positive pools are considered, and their list of specimens is updated such that specimens that appeared in at least one negative pool are ‘eliminated’. An optimal elimination results in at least one positive pool that contains a single specimen 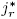, which is retrieved as the detected specimen that contains *r*, and the algorithm halts. However, in case that following elimination all positive pools still contain several specimens, the next step is used to retrieve the most probable specimen that contains *r*.
- *Likelihood step:* Following Elimination each of the *T* ^+^ positive pools contains 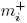 specimens, where *i* = 1,..,*T*^+^ (the smallest possible size of 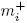 is 2 otherwise an specimen would have already been selected in the former Elimination step). We aim to retrieve an specimen 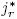 that maximizes the likelihood of the measurement vector *y_r_*: 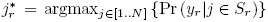. Using Bayes law and assuming that the prior probability Pr (*j* ∈ *S_r_*) is equal for all specimens, then 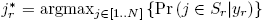. Since calculating Pr (*j* ∈ *S_r_*|*y_r_*) is computationally hard, we use the following approximation. Recall that every positive pool must contain at least one specimen from *S_r_*. Intuitively, an specimen that belongs to many small positive pools should be preferred over an specimen that belongs to a few large positive pools. Hence we calculate the probability of an specimen to be chosen when uniformly and independently selecting one specimen from each positive pool. The specimen *ĵ*\* with the highest probability of being selected at least once is retrieved, 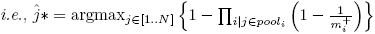.

### Measurement matrix design

The measurement matrix *M* should be designed so as to support the detection algorithm, *i.e*., to increase the instances in which the Elimination step suffices for detection, and provide correct detection by the Likelihood step when applied. Intuitively, this calls for a ‘continuum’ of pool sizes, where small pools are aimed to detect the highly abundant specimens while larger pools would ‘catch’ the less abundant ones. In addition, *M* should reflect common experimental constraints, *e.g*., it should keep a low number of specimens per pool and a low number of pools that contain each specimen. The latter reflects a limit in material while the former stems from sequencing considerations since PCR amplification and sequencing of very large pools may introduce noise due to, *e.g*., PCR biases.

In case there is *exactly* one positive specimen *j* out of the *N* specimens, the algorithm would almost always correctly detect it for any designed matrix *M*, assuming its columns are unique, since all specimens except *j* would be eliminated. However, correct detection of specimens whose abundance is higher than 1/*N* requires carefully designing *M*. Intuitively, the design aims to increase the probability of creating pools that contain exactly one positive specimen, irrespective of the specimen’s abundance. In such cases the positive specimen can be detected with high probability.

The design is comprised of five steps (Fig 3). The first step randomly splits the specimens into non-overlapping sets termed *levels*, whose sizes form an increasing geometric series. Since the expected number of positive specimens in a level also forms an increasing geometric series starting from size 2, there exists a level in which the expected number of positive specimens is *about* 1. Second, a certain number of pools is allocated to each level, *i.e*., these pools would eventually contain specimens only from their corresponding level. The first two steps are illustrated by the black blocks in Fig 3A. In the third step the blocks are ‘smoothed’ by interpolating the top and bottom pools of each specimen resulting by the upper and lower black curves in Fig 3B. The fourth step randomly assigns specimens to pools according to these limits, *i.e*, each specimen may only be included in the subset of pools determined by the upper and lower limits. The fifth step verifies that the columns of the resulting matrix *M* are unique, and repeats the allocation of specimens to pools for columns, *i.e*., specimens, that were found to be non unique. The following sections present details of the first four steps.

**Figure 3:**
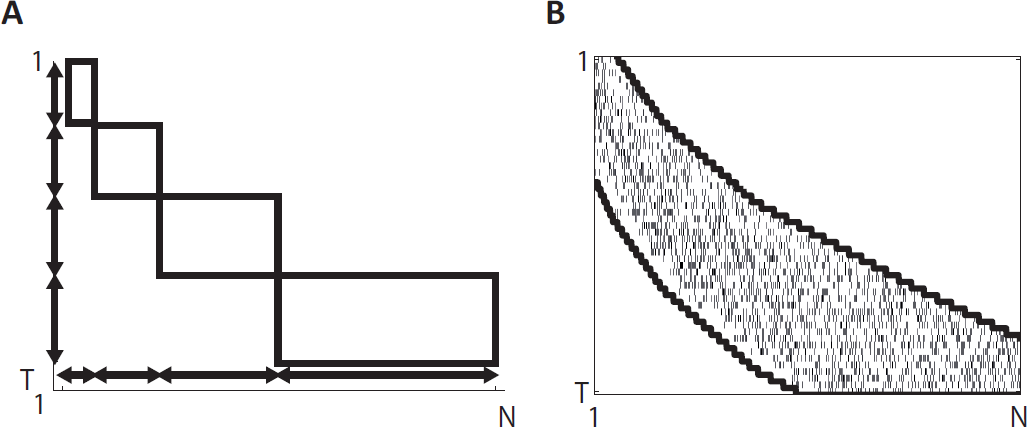
Designing the measurement matrix *M*. A. The first two steps of the construction. Specimens are first divided into *levels* shown as horizontal double headed arrows (since the specimens’ order is random so is the assignment). Second, pools are allocated to each level, as shown by vertical arrows. The first two steps result in the black blocks. At this stage specimens may only be assigned to pools according to these blocks. B. ‘Smoothing’ the blocks by interpolations and actual assignment of specimens to pools. Smoothing results in the upper and lower curves. Each specimen is then randomly assigned to a subset of the pools between the upper and lower limits (a small tick denotes an allocation of an specimen to a pool).

#### Dividing specimens into levels

The design should support any abundance between *a_min_* and *a_max_*. The latter was set to 0.5 and the former is effectively 2/*N*, since the lowest possible abundance 1/*N* is supported due to the uniqueness of the columns of *M*. Let *L* be the number of levels, and *N_l_* be the number of specimens in the *l*’th level. We set *N*_1_ such that the expected number of positive specimens at the top level is 1 for *a_max_, i.e., N*_1_ = 2, and similarly for the bottom level *N_L_* =1/*a_min_* = *N/*2. Since levels’ sizes increase geometrically with a ratio *q*, the size of the *L*’th level is *N_L_* = *N*_1_ · *q*^*L*–1^, and the sum of all levels’ sizes is 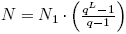, which allows solving for *q* and *L*.

#### Allocating pools to each level

In case there is exactly one positive specimen in level *l*, then log_2_ (*N_l_*) pools are required for detection. Since the sizes of the levels form a geometric series, log_2_ of the size of the levels forms an arithmetic series. Let *T_l_* = *c* log_2_ (*N_l_*) be the total number of pools at level *l*, where *c* is a proportion constant, which can be calculated using the equation for the sum of an arithmetic series, 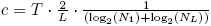. Dividing pools into levels followed by allocating pools to each level results in the blocks displayed in Fig 3A.

#### ‘Smoothing’ - Interpolating the blocks

Although there exists a level in which the *expected* number of positive specimens in each level is *about* one, it may happen that no level contains *exactly* one positive specimen. There may exist adjacent levels (*l, l* + 1) that display a transition between zero and two or more positive specimens. We aim to ‘smooth’ this transition, *i.e*., create pools that mix specimens from levels *l* and *l* + 1, so as to increase the probability of creating a pool with a single positive specimen (levels that do not contain any positive specimen and levels that are flooded with many positive specimens are useless with respect to detecting a positive specimen). Smoothing results in the upper and lower limits in Fig 3B. The rationale for smoothing is described, while the exact details of the heuristic used appear in the Supplementary Methods.

The rationale for creating pools that mix level *l* +1 specimens with level *l* specimens is two fold. First, in case there are zero positive specimens in level *l* and several positive specimens in level *l* + 1, then the pools of level *l* are wasted. Creating pools that add *part* of level’s *l* +1 specimens to level *l*, increases the probability of having a single positive specimen in the modified pools. Second, to increase the number of eliminated negative specimens in level *l*, mixing allows specimens from level *l* to also participate in pools allocated to level *l* + 1, and hence in case such pools turn out to be negative, the specimens that belong to level *l* are eliminated. On the other hand, if a pool from level *l* + 1 contains one or more positive specimens then adding positive specimens from level *l* does not significantly reduce performance since the likelihood objective function is mostly influenced by the smaller pools of level *l*.

#### Assigning specimens to pools

Each specimen is randomly assigned to a fraction *α* of the pools defined by the upper and lower limit for that specimen. The value of *α* is selected by trial and error; for each value of *N* and *T* random realizations of *M* are created for different values of *α* and the best one is selected (typically 0.2 < *α* < 0.3).

*Remark*s: The above design follows experimental considerations that constrain both the pool size and the number of pools in which each specimen participates. Most specimens participate in approximately the same number of pools, which also eliminates an unwanted bias in the likelihood selection step.

### Example

As an introductory example we present the performance of a specific design for *N* = 1000 cultures and *T* = 84 pools. A matrix *M* was created as described, and detection was tested in the following way. For each abundance in the range from 1/1000 to 500/1000 we randomly selected a subset of positive specimens and calculated the corresponding measurement vector *y*. Detection was applied and success probability was estimated based on 10^5^ simulations for each abundance (Fig 4). Success probability is higher than 95% for the whole range of 0% – 50%. By construction, the design detects the lowest abundance of 1/1000 with 100% success, due to the uniqueness of the columns of *M* and in the absence of measurement noise. The effects of each of the two algorithmic steps in GT-Census are shown in Supplementary Methods.

**Figure 4:**
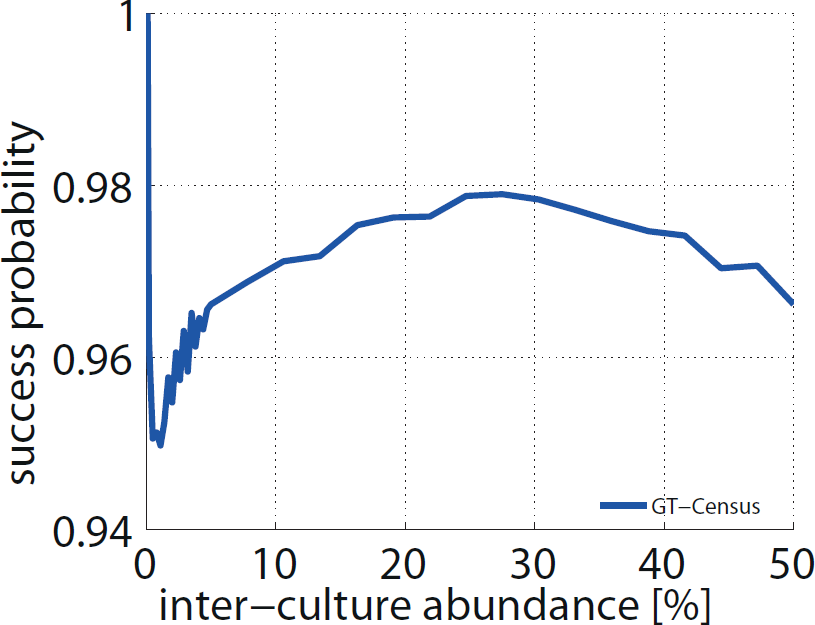
Success probability - Introductory example. Success probability in detecting a positive specimen out of 1000 specimens using 84 pools in the noiseless case, as a function of inter-culture abundance.

### Experimental noise

Several types of experimental noise factors should be taken into account when designing GT-Census pooling experiments, as outlined below.

#### Overcoming dropped pools

Sequencing experiments may include dropout pools, namely pools that failed to be amplified and result in zero reads. This type of noise is easily detected, and redundant pools should be added to overcome it, as shown in the Results.

#### Overcoming false positive callings

A pool may be called positive for *r* even if the bacterium was not actually in the pool. Such false positive calling may be caused by cross-pool contamination or by read errors (*e.g*., a highly similar bacterium *r′* appears in the pool and read errors cause its reads to be mapped to *r*). False positive callings can be alleviated by increasing the threshold required for calling positive pools.

#### Overcoming false negative callings using a known spike

False negative callings, *i.e*., mis-calling specific bacteria, may be caused by the following factors:

a. One or more bacteria ‘overtake’ a pool: To avoid such a scenario equal amounts of DNA should be taken from each specimen. This problem may be more relevant in the scenario of cultures. To alleviate the problem all cultures should be allowed to grow until saturated, so as to overcome different growth rates among bacteria. Measuring DNA concentrations of each culture or sample overcomes this problem, although being a costly stage that may be avoided.
b. Insufficient number of reads: A low number of reads per pool may result in mis-called bacteria, especially in pools that contain low abundance bacteria. Reads should be allocated in proportion to the number of specimens per pool (in the case of cultures) and also according to the smallest required intra-sample abundance (in the case of samples). This can be done by scaling the amount of sequenced material by the pool’s size.
c. PCR bias: PCR amplification of a pool may cause biases, especially in large pools, and hence bacteria may ‘disappear’. It has been shown that almost uniform amplification can be reached when pooling more than 500 samples [24, 25] which makes PCR bias less of a problem in our design. To reduce potential PCR bias the GT-Census pooling design limits pool size.

Taking the relevant precautions described in (a)-(c) the number of false negative callings should be minimized. However, to further deal with false negatives, we suggest to spike a known amount of a specific bacterium to each pool. Such a positive control would provide an estimate of the probability of a false negative calling, *p_fn_*, and allow a modification in the detection algorithm that accounts for such mis-callings.

### Detection algorithm in the presence of noise

False negative measurements require a modified detection algorithm, since strict elimination can not be used (a pool may be negative due to mis-calling and hence elimination of the pool’s specimens may be incorrect). Hence, instead specimens are stochastically eliminated, followed by a modified likelihood step. Moreover, as observed empirically, the cases of cultures and of samples require different detection algorithms as described below.

#### Detection in a repository of *cultures*

The following two step algorithm is used, assuming that the false negative calling, *p_fn_*, was experimentally estimated:

- *Stochastic Elimination step*: In the absence of measurement noise, an specimen *j* was eliminated if it appeared in at least one negative pool, and was retained if it appeared only in positive pools. To accommodate for false negative calls, we define *w_j_*, the retention probability of specimen *j*. Assuming that false negative calls are independent across pools, then 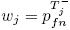, where 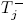 is the number of negative pools of specimen *j*. Hence, specimens are eliminated with probability 1 – *w_j_*, and the following Likelihood step is applied.
- *Likelihood step*: As in the noiseless case the specimen 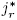 with the highest likelihood of being positive, 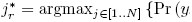 should be retrieved. Using an analogous argument Pr (*j* ∈ *S_r_*|*y_r_*) is approximated by blackthe probability of an specimen to be selected from each positive pool. The probability of selecting specimen *j* in pool *i* is equal to 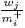, where 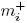 is the sum of the retention probabilities of specimens in the positive pool *i*. The retrieved specimen, *ĵ*\* maximizes the objective function 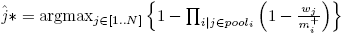.

*Remark*: In the absence of noise, *i.e*, when *p_fn_* = 0, the algorithm is identical to the one described for the noiseless case (an specimen observed in at least one negative pool 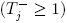 is eliminated since 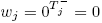, while otherwise *w_j_* = 1).

#### Detection in a repository of *samples*

When mining a repository of monomicrobial cultures all occurrences of bacterium *r* are symmetric, and hence any retrieved culture that contains *r* suffices. In contrast, when mining a repository of samples it is desirable to retrieve a sample in which the abundance of *r* is high enough. To allow this, the selected specimen, *ĵ*\* maximizes the modified objective function 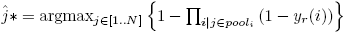, where *y_r_*(*i*) is the fraction of reads of bacterium *r* in pool *i*.

### Simulations

Extensive *in silico* simulations were performed to evaluate the performance of GT-Census, as described separately for the case of censusing repositories of cultures or of samples.

#### Censusing a repository of *cultures*

For each pair of *T* and *N* a matrix *M* was created, as described in ‘Measurement matrix design’. Simulations were performed for abundances in the range 0% – 50%, where for each abundance *a, a · N* positive specimens were randomly selected and the resulting measurement vector *y* was created. When simulating a false negative classification probability *p_fn_*, each non-zero entry in *y* had a probability *p_fn_* of being zeroed. Simulating dropout pools, a certain fraction of the pools was randomly selected and deleted. Given the measurement vectors *y* and the matrix *M*, the detection algorithm was applied and the correctness of the retrieved representative was checked. This procedure was repeated 10^5^ times for each value of *a*. A matrix *M* was termed ‘valid’ for a pair (*N, T*) if the detection success rate was higher than 0.95 for all values of *a* in the range 0% – 50%.

#### Censusing a repository of *samples*

Censusing a repository of samples depends on intra-sample bacterial abundance. To apply representative intra-sample abundances we simulated two GT-Census experiments using reads from large scale experimental projects, namely the American Gut project [22] and the household data of Lax et al. [23].

#### Simulating censusing a repository of *samples* - the American Gut experimental data

The American Gut project sequenced samples from several body sites of more that 10, 000 American volunteers. We randomly selected a subset of 1, 000 fecal samples and simulated a GT-Census ‘experiment’.

*The naive approach - creating the ground truth*: For each of the 1000 selected samples the list of OTUs was created by the QIIME package [26], using raw FASTQ files (downloaded from the QIITA server, study ID 10317). The number of reads per sample varied between 14, 000 – 426, 000 and were used as follows (reads were not rarefied). Low quality reads were discarded followed by closed-reference OTU picking (at levels of 97% and 99% identity). OTUs whose intra-sample abundance was higher than 1% were subsequently considered to create the ground truth list of OTUs across the 1, 000 samples, resulting in 1344 and 1663 OTUs for 97% and 99% identity, respectively. Figure 5A shows the distribution of 97%-OTU abundance across 1, 000 samples. Most OTUs are rare, which makes applying GT-Census highly attractive in this case.

**Figure 5:**
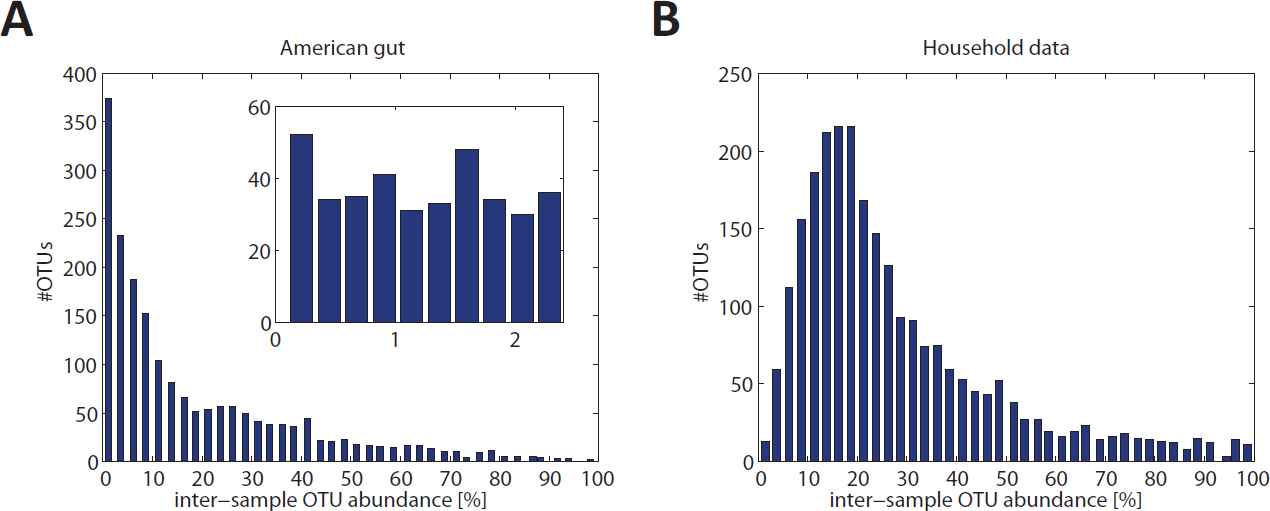
Abundance of OTUs across samples. A. Abundance of OTUs (at a level of 97% similarity) across samples for American Gut data. The inset shows a finer resolution of the lowest histogram bin. B. The same for the household dataset of Lax et al.

*‘Sequencing’ a pool*: A GT-Census design was created for 1, 000 samples comprising of 100 pools. The design was set to tolerate up to 1% false negative detections. To create a pooled ‘experiment’ either 10, 000 or 1, 000 reads were randomly selected from each sample that participated in a pool (reads were randomly selected directly from the sample’s FASTQ file without prior filtering for high quality reads).

*Analyzing GT-Census results*: To allow for a comparison between the naive approach and GT-Census, QIIME was applied to each of the pools in same way as described for individual samples, creating a list of OTUs in each pool. The intra-pool fraction of reads mapped to each OTU was also recorded. Since we were interested in censusing OTUs for which the abundance of the OTU was higher than 1%, the list of potential OTUs was created in the following way. OTUs for which the intra-pool fraction of reads times the pool’s size was higher than 1% were considered. This is a necessary condition for an intra-sample abundance of more than 1%, although it may happen that GT-Census was also applied to OTUs whose actual intra-sample abundance is lower (*e.g*., two samples that contain an OTU in abundance 0.5% are pooled and hence the effective OTU abundance is 1%). GT-Census was applied to each of these OTUs resulting in a predicted sample for each OTU. These predictions were then validated by the ground truth and the percentage of correct GT-census calls was calculated.

#### Simulating censusing a repository of *samples* - Household data

Lax *et al*. [23] collected samples from seven US families and their homes over six weeks, comprising of 1625 samples whose V4 region were sequenced using an Illumina machine. The authors used these data to study the correlation between the house microbiota and the family that lived in it. We used this dataset to display the potential advantages of GT-Census in detecting all bacterial species across these different niches. Detection correctness was compared to the naive approach of sequencing each of the samples individually, as described for the American Gut dataset.

*The naive approach - creating the ground truth*: Samples with less than 10, 000 reads were ignored. For each of the 1559 remaining samples the list of OTUs (at the level of 97% similarity) was created using the raw FASTQ files (downloaded from the QIITA server, study ID 2192). The number of reads per sample varied between 37, 000 – 680, 000. The total number of OTUs across samples was 1420. Since samples were collected along time and in adjacent places (*e.g*., a house’s bedroom and bathroom along time), it was expected that microbial content would display larger overlap across samples than in the case of the American Gut dataset (Fig 5B).

*‘Sequencing’ a pool and analysis of GT-Census results via QIIME*: A GT-Census design was created for 1559 samples comprising of 155 pools, which allows to overcome up to 1% false negative detections. Pooled ‘sequencing’ and performing GT-Census over QIIME results was performed in the same way as described for the American Gut dataset.

## 3 RESULTS

The performance of the GT-Census approach was evaluated by extensive *in silico* simulations. GT-Census of cultures was initially considered since it does not involve assumptions regarding intra-sample bacterial abundance, and thus allows to display different aspect of the approach.

GT-Census of *cultures*: The noiseless case was considered providing an estimate for the minimal number of pools required for a design to be ‘valid’, *i.e*., to achieve 95% accuracy for all abundances in the range 0%–50%. Second, the effect of experimental errors on the required number of pools was evaluated. Third, the number of reads required as a function of the number of cultures was evaluated, to serve as a proxy to sequencing costs.

GT-Census of *samples*: We show example applications of GT-Census based on experimental data from the American Gut project [22] and from the household longitudinal study data [23]. These examples display different intra-sample and inter-sample bacterial abundances and allow the evaluation of GT-Census in two realistic scenarios.

The code used for the simulation is available at https://github.com/NoamShental/GT-Census.

### Cultures - Number of required pools and code efficiency in the noiseless case

Fig 6A presents the minimal number of pools required for a ‘valid’ construction, *T_min_*, as a function of the number of cultures *N* in the absence of experimental noise. Such *T_min_* assures detection success in at least 95% of the cases for all abundances in the range 0% – 50% (0% corresponds to an abundance of 1/*N*). Fig 6B presents the gained efficiency in those settings, *i.e*., the ratio *N/T_min_*. For example, for *N* = 5000 cultures our construction required about 130 pools, corresponding to more than 35 fold decrease in required resources compared to the naive approach.

**Figure 6:**
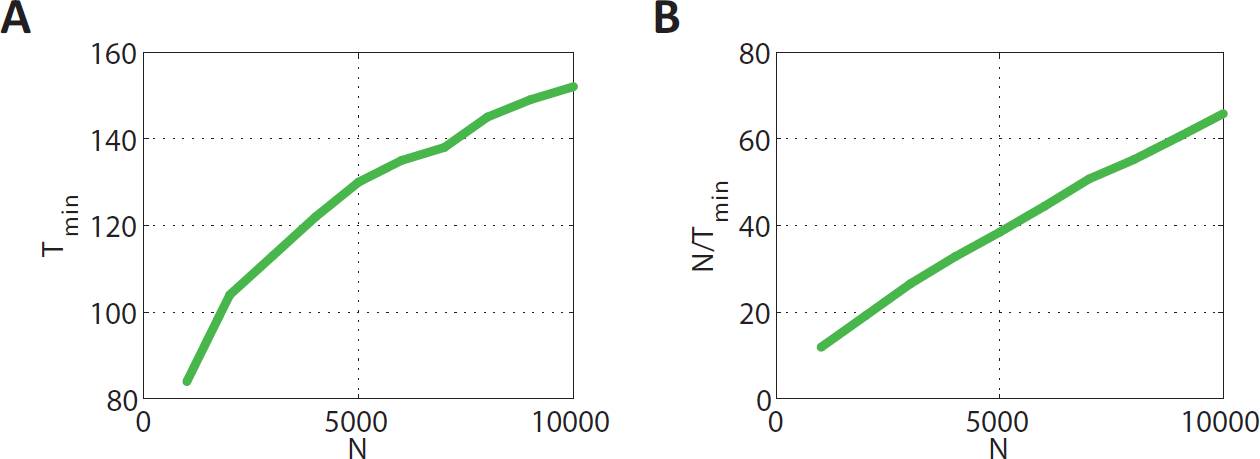
GT-Census Performance in the noiseless case. A. The minimal number of pools, *T_min_*, required for a ‘valid’ design that supports all bacterial species having an abundance of 0% – 50% as a function of the number of cultures *N*. In a ‘valid’ pair of *N* and *T_min_* at least 95% of the simulations successfully identified a positive culture for each abundance in the range. B. The efficiency score, namely *N/T_min_* for the setting described in (A), corresponding to the reduction in required resources in fold change.

As expected, *T_min_* increased as *N* increases, however the efficiency score *N/T_min_* also increased, namely applying GT-Census becomes more cost effective when increasing the number of cultures (since the number of pools allocated to each level is *O* (log (*N*)) and the number of levels is *O* (log (*N*)), the total number of pools is *O* log^2^ (*N*)), and hence the efficiency ratio is *O*(*N/* log^2^(*N*)) which grows with *N*).

### Cultures - Robustness to experimental noise

Two types of potential experimental measurement errors are considered. Fig 7A shows *T_min_* as a function of *N* in case 1% or 5% of the pools are randomly deleted. The value of *T_min_* increases compared to the noiseless case (green line) by a factor close to the percentage of deleted pools. Hence, an experimental design should increase the number of pools roughly by the expected dropout rate.

**Figure 7:**
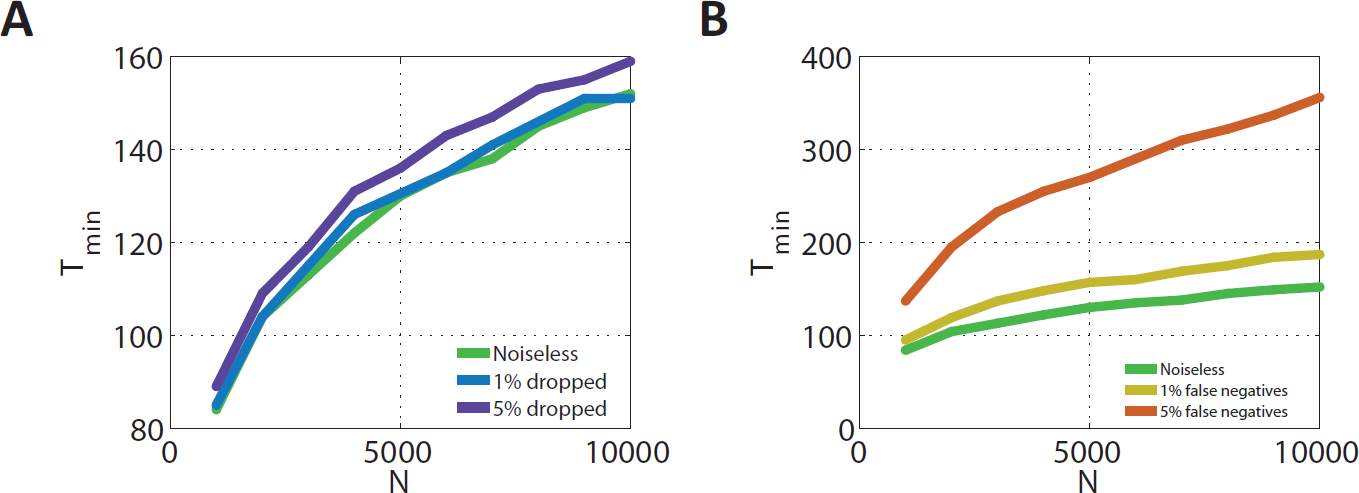
Robustness to experimental noise. The minimal number of pools required for a ‘valid’ design as a function of the number of cultures for a certain percentage of dropout pools (A) or false negative classifications (B). The noiseless case appears in green.

Fig 7B shows *T_min_* as a function of *N* in case of false negative calls. Each positive pool has a probability of 1% and 5% to be falsely classified as negative (the pool itself is not dropped namely it may be positive for other bacteria). The effect of false negative detections is more pronounced, which results in an increase in *T_min_*, compared to the noiseless case (green line) especially for 5% false negatives. However, efficiency remains high even in the presence of false negative detections.

### Cultures - An estimate of the required sequencing resources

Figures 8A–B show the total size of pools as a function of the number of cultures for the designs in Figs 7(A,B), respectively. The total number of cultures across pools grows almost linearly with *N*, and thus serves as a proxy for the required sequencing costs. For example, screening a set of 10, 000 cultures required sampling about 90, 000 cultures across all pools. When censusing a set of 10, 000 cultures, and assuming that 100 reads are required for calling a positive pool for a certain bacterium, then the total number of reads is less than ten million, taking only a fraction of a standard Illumina lane. Censusing a set of 10, 000 samples, and assuming that the lowest relevant intra-sample abundance of a bacterium is 1%, requires about 10^9^ reads, *i.e*., about one Illumina flow cell or one lane of the new Illumina machines.

**Figure 8:**
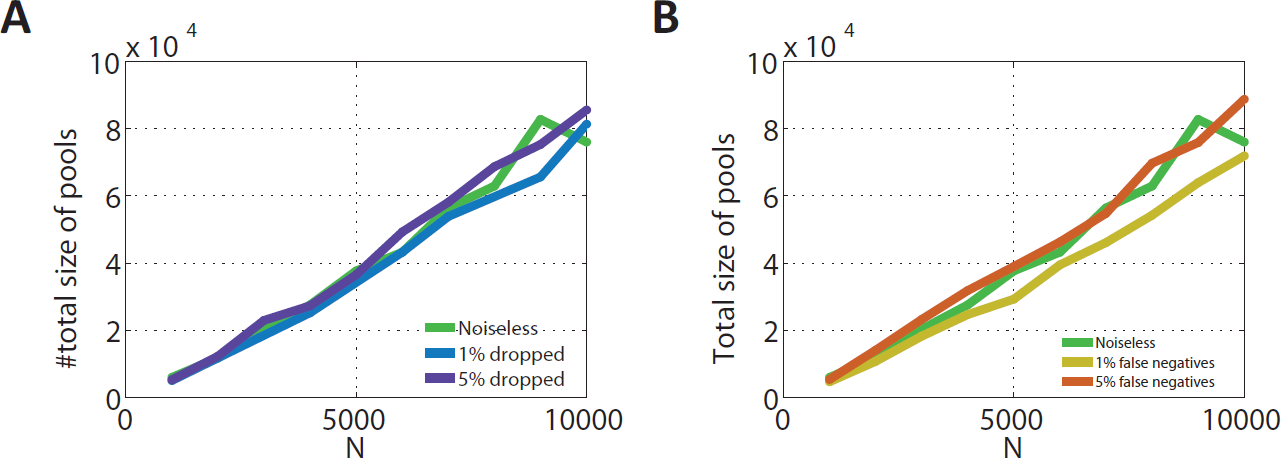
The total pools’ sizes. The total number of cultures across all pools for the cases appearing in Figures 7(A,B), respectively. This number serves as a proxy for the required number of reads and hence to sequencing costs.

### Samples - High performance of GT-Census in a simulated experiment using large scale projects

To evaluate the potential performance of GT-Census over real samples we performed two independent tests over the American Gut project and the household data of Lax *et al*. The ground truth for each dataset was created by the list of OTUs that appeared in each sample (OTUs whose intra-sample abundance were higher than 1% were considered). A GT-Census ‘experiment’ was performed for each dataset, measuring the probability of correctly detecting a sample that contains each of the OTUs that appear across samples as a function of inter-sample OTU abundance (Fig 9). Performance was higher than 90% for all inter-sample abundances in both datasets and there were no additional false positive OTUs detected.

**Figure 9:**
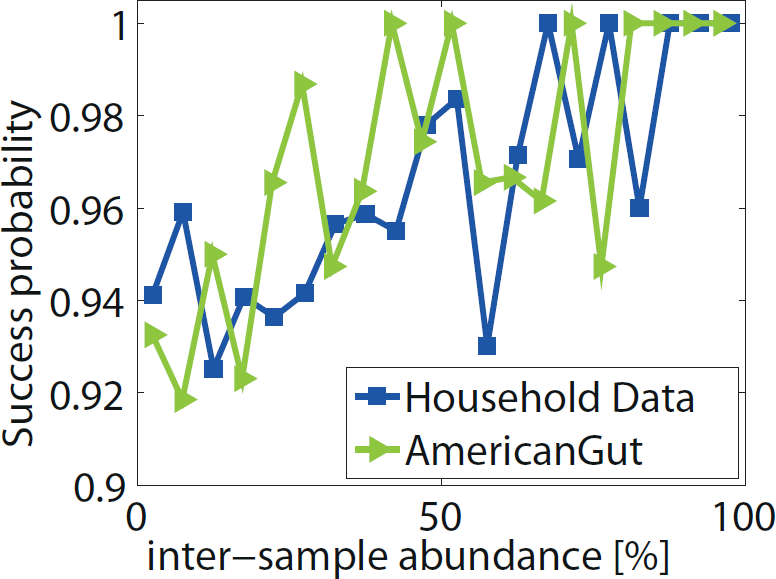
GT-Census performance over the American Gut and the household data of Lax et al. Probability of correctly detecting a sample that contains an OTU as a function of inter-sample OTU abundance for two independent tests of GT-Census.

These examples display the potential of applying GT-Census for detecting a sample for each and every bacterial species within a large cohort of samples using only a fraction of resources. The number of pools was 100 and 155, for the American Gut and household data, respectively, corresponding to a ten fold saving in resources compared to the naive approach. The total number of reads allocated across all pools was 7 10^7^ and 1.7 · 10^8^, for the American Gut and household data, respectively. Hence GT-Census may enable highly efficient censusing using a ten fold less costly and labor intensive sample preparation, while taking a smaller number of reads than were used in the original experiments (performance for a lower number of reads and other comparisons displaying the robustness of GT-Census appear in the Supplementary Results).

## 4 DISCUSSION

GT-Census presents a novel approach for bacterial diversity censusing of large repositories of either cultures or samples. Combining sequencing and group testing provides high resolution censusing for only a small fraction of costs required by the naive approach. Specifically, for the case of censusing bacteria in a repository of samples, GT-Census provides the first affordable solution. Although we tried to take into consideration various noise effects and simulated pools from experimental reads of two large scale studies, experimental validation of GT-Census is required to evaluate its performance.

*Potential limitation of GT-Census*: Scaleability of GT-Census to very large repositories is limited by the number of specimens that can be pooled together so as to avoid false negative measurements. A uniform amplification of pools of more than 500 samples was achieved in several studies [24, 25] yet larger pools need to be tested.

*GT-Census in the lens of group testing*: Computationally, GT-Census presents a novel group testing problem that calls for rigorous theoretical analysis which is beyond the scope of this manuscript. Such analysis would improve the pooling design and the detection algorithm presented in this manuscript. Also, this problem calls for rigorous performance analysis that would set theoretical bounds on the lowest possible number of pools required in a given setting.

*Variations of GT-Census*: Many more potential variations of GT-Census may be interesting to follow in the future. For example, when censusing a repository of samples it might be interesting to modify the objective such that to detect a representative sample that contains a certain *set* of bacteria, instead of a *single* bacterium. For example, one can seek a representative sample that contains both bacteria “A and B” or “A and B but not bacterium C”. All such queries can be performed *a posteriori*, hence answering various queries. The current pooling matrix is not optimized for complex queries, and it would be interesting to design such matrices (the experimental part of GT-Census would remain the same).

*Other applications of GT-Census*: GT-Census was presented in the context of censusing bacteria using NGS. Another interesting direction may be to apply GT-Census via single-cell genomics, currently used for whole genome sequencing of single bacterial cells [27]. Such initiatives involve repeated whole genome sequencing of the same cells, which may be replaced by GT-Census. A second potential direction may be censusing of *genes* instead of organisms. For example it might be interesting to detect samples that contain “gene A and gene B, but not gene C”. Analogously, GT-Census may be applied to census meta-transcriptomic, meta-proteomic and metabolomic data. The latter two applications involves applying GT-Census over Mass Spectrometry data, which may become highly useful in the future. Third, GT-Census may also be applied in different contexts, other than in microbiology related applications. This solution-in-search-of-a-problem computational approach is suitable whenever seeking a representative specimen for each potential phenotype out of a repository of unknown specimens, *e.g*., screening large drug libraries.

## 5 ACKNOWLEDGEMENTS

We thank Tal Luzzatto-Knaan for many useful discussions.

## Shalem et al. Supplementary Material

### 1 Supplementary Methods

#### 1.1 Detailed measurement matrix design

Items are divided into levels, and each level is allocated a number of pools (Figure S1A). The next step interpolates the blocks such that pools’ sizes gradually increase (Figure S1C). Interpolation is described below.

*Interpolation of the upper limit*: The first *n_i_* items of level *i* + 1 are added to the *n_i_* items of level *i* by connecting the interpolation points (red points in Figure S1B).

*Interpolation of the lower limit*: The lower limit is set at a constant offset from the upper limit until we reach the last pool, *T* (and from that point on it equals the last pool). The offset’s size (yellow arrow in FigureS1B) is set by the distance between the total number of pools *T* and the upper limit of the first item of the bottom level. The lower limit essentially increases the number of pools allocated to level *i*.

**Figure S1:**
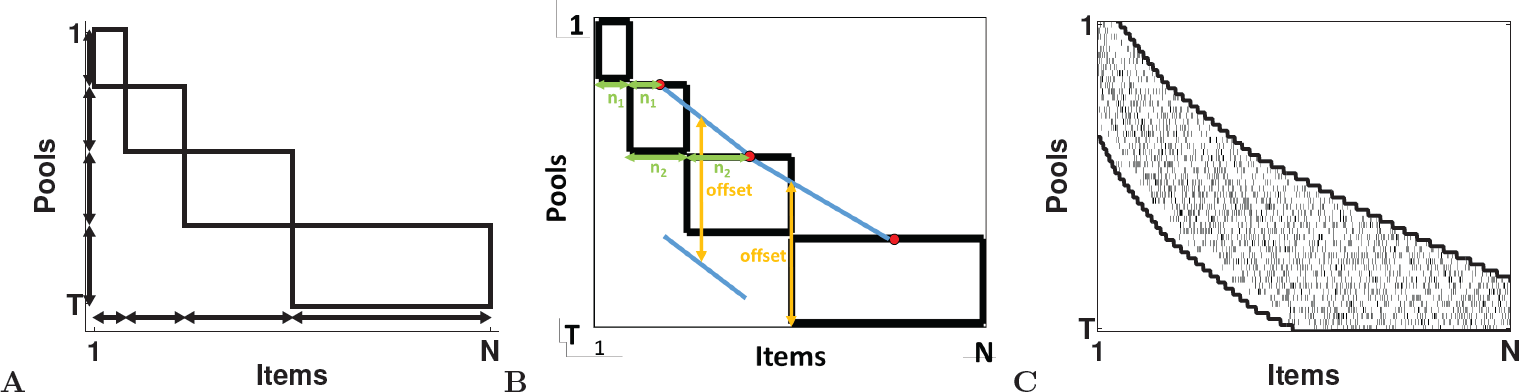
Designing the measurement matrix. A. The first step of the construction. Items are first assigned to levels (shown as horizontal double-headed arrows), and then pools are assigned to each level (shown as vertical double-headed arrows arrows), resulting in the black rectangles. B. Smoothing the rectangles by interpolations. C. The resulting upper and lower limits. Each item is randomly assigned to a subset of the pools between the upper and lower limits (a small tick appears whenever an item is allocated to a pool).

#### 1.2 Example - the detection algorithm

Materials and Methods presents an introductory example of the detection algorithm in the noiseless case. In brief, the performance of a specific design for *N* = 1000 cultures and *T* = 84 pools was evaluated (Figure 4 and blue line in Figure S2). For each abundance in the range from 1/1000 to 500/1000 a subset of positive items were randomly selected, detection was applied and success probability was estimated based on 10^5^ simulations. Success probability is higher than 95% for the whole range of 0% – 50%.

GT-Census detection algorithm is comprised of an Elimination step, followed by a Likelihood step whenever Elimination does not suffice. This section uses the introductory example to study each of these steps.

*Elimination-only algorithm*: A ‘good’ matrix *M* should allow Elimination in most cases and only rarely require using the Likelihood step. The black line in Figure S2 presents the success probability when only the Elimination step was required, namely the fraction of cases where elimination created a pool with a single positive item. In about 85 – 90% of the cases *M* supports full elimination.

*Elimination followed by random selection algorithm:* The red line in Figure S2 presents the performance when applying the Elimination step but replacing the Likelihood step by randomly selecting one item from the smallest positive pool, in case Elimination did not suffice. Hence applying the Likelihood step gains about 5% in performance (*i.e*., the difference between the blue and red lines).

**Figure S2:**
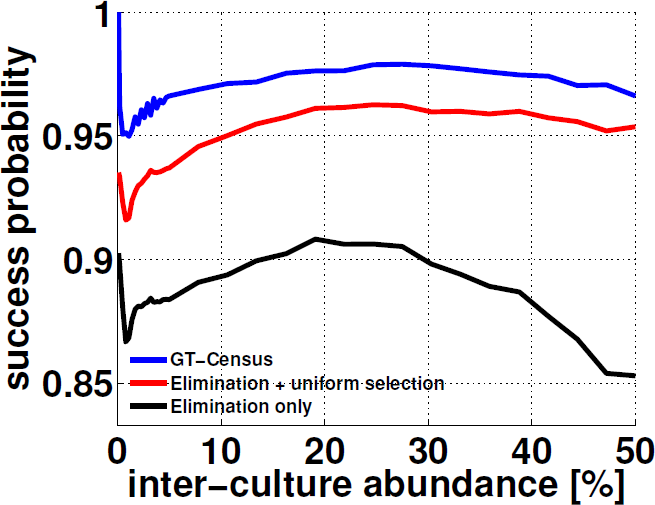
success probability vs. abundance. Success probability of GT-Census in detecting a positive item out of 1000 items using 84 pools in the noiseless case, as a function of inter-culture abundance (blue curve). The black curve shows performance in case only the Elimination step of GT-Census is applied. The red curve corresponds to using Elimination followed by randomly selecting an item from the smallest positive pool, instead of the maximum likelihood approximation.

### 2 Supplementary Results

#### 2.1 Simulating a GT-Census experiment using the American Gut project data

Figures S3A–C show the performance of GT-Census in censusing the American Gut pooling ‘experiment’. Each subplot presents the former results of the ‘standard’ setting (main text Fig 9, and green line), and the performance of GT-Census when modifying one of these parameters. In the ‘standard’ setting detection was performed using the *sample* ‘version’ of GT-Census; the number of reads per sample in each pool was 10, 000; the performance was tested over detecting OTUs at a 97% identity level; and the threshold for intra-pool abundance was 1%.

*The effect of using the Detection algorithm of cultures*: The samples ‘version’ of the GT-Census Detection algorithm uses the fraction of reads assigned to an OTU in a pool, rather than the binary absent-present calling applied for *cultures*. Using the latter as the GT-Census detection algorithm decreased performance by up to 10% (Figure S3A) displaying the importance of using the measurements’ continuous values as rather than simple binary absent-present calls.

*The effect of varying the number of reads*: Reducing the number of reads per sample in a pool, from 10, 000 to 1, 000 lowered GT-Census performance down to 80% success for low abundance OTUs. OTUs whose intra-sample abundance was around 1% were sometimes missed altogether when allocating 1, 000 reads per sample, and hence performance was lower.

*The effect of changing the required phylogenetic resolution*: We repeated the whole simulation using an OTU level of 99% identity instead of the 97% level. QIIME was applied to all individual samples at a level of 99% to set the ground truth, resulting in 1663 OTUs across samples. GT-Census was then applied to the same set of pools as in the former case, while considering OTUs at a level of 99%. Although the number of OTUs increased compared to the level of 97%, success probability of GT-Census did not vary, *i.e*., higher phylogenetic resolution may be sought with no apparent decrease in performance.

*The effect of seeking OTUs of lower intra-sample abundance*: The simulation was applied to OTUs whose intra-pool abundance times the pool size was higher than 1% (this served as a proxy to the intra-sample abundance of 1%). The number of reads in the standard setting was 10, 000 per sample in a pool, resulting in more than 90% success rate for all OTUs. The threshold of 1% was varied to test the robustness of results (Figure S3D). Performance remained higher than 90% up to a threshold of 0.5%. For a threshold of 0.05% success probability decreased to a level of 80% (for low inter-sample abundances), and results went down to 70% for a threshold of 0.01%. Hence, the threshold for the lowest required intra-sample OTU abundance should be roughly correlated to the number of reads allocated.

**Figure S3:**
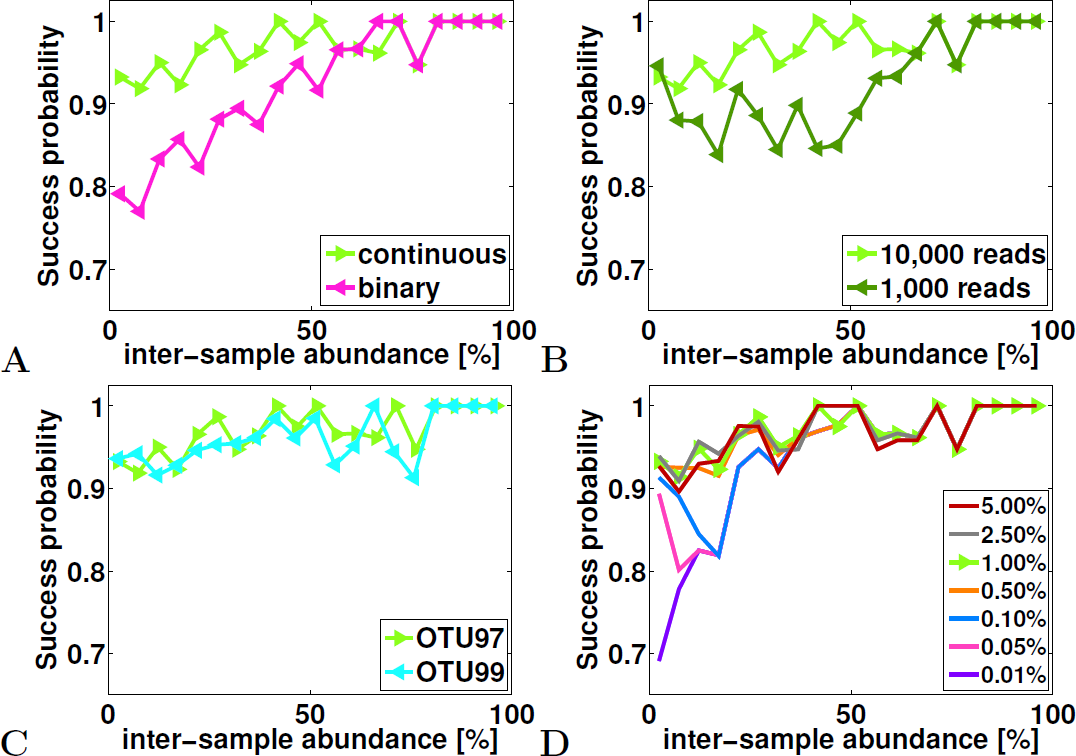
GT-Census for the American Gut data set. Success probability of GT-Census as a function of inter-sample OTU abundance. The green line in each subplot corresponds to the standard GT-Census setting, while modifying the GT-Census Detection algorithm (A), reducing the number of reads (B), seeking OTUs of 99% similarity (C), and for lowering the threshold for required intra-sample OTU abundance (D).

